# *In silico* assessment of missense point mutations on human cathelicidin LL-37

**DOI:** 10.1101/2021.01.30.428671

**Authors:** William F. Porto, Sergio A. Alencar

## Abstract

Cathelicidin antimicrobial peptides are a diverse family of cationic amphipathic peptides with multiple activities. In humans, cathelicidin LL-37 is one of the main host defense peptides with a remarkable medical and biotechnological potential. Deregulation of LL-37 expression has been associated with inflammatory diseases. However the effects of point mutations driven by single nucleotide polymorphisms (SNPs) on LL-37 are unknown. Here we applied an array of computational tools to investigate the effects of such mutations on LL-37 structure and activity. Due to the fact that, on cathelicidins, the prodomain is more conserved than the mature peptide, the SNP effect predictions were biased and, overall, resulted in neutral effects; and due to the slight changes in physicochemical properties, the antimicrobial predictions indicated the maintenance of such activity. Nonetheless, R07P, R07W, R29Q, R29W mutations reduced the peptide net charge, which in turn could result in less active LL-37 variants. Molecular dynamics data indicated that R07Q and N30Y mutations altered the LL-37 structure, leading to potential deleterious effects. In addition, the helix dipole is altered in G03A, R07P, R07W and L31P mutations, which could also alter the antimicrobial activity. Our results indicated that despite the mutations did not alter the residues from LL-37 active core, they could influence the antimicrobial activity and consequently, could be involved in inflammatory diseases.

## 1 Introduction

Antimicrobial peptides (AMPs) have been proposed as promising alternatives to conventional antibiotics. AMPs are usually short, cationic, amphipathic peptides found as components of innate immune system [1,2]. Cathelicidin AMPs are a diverse family of AMPs varying from 12 to 100 amino acid residues [3]. At first look, cathelicidins compose a very heterogeneous group. However, they are grouped together due to their precursor sequence, which has a cathelin domain [3,4]. In many vertebrates, including humans, the cathelicidins, together with defensins, represent a large group of host defense peptides (HDPs) [1,4].

In humans, *CAMP* gene, located at 3p21.31, encodes the cathelicidin antimicrobial peptide LL-37 as a precursor protein, containing the cathelin domain and the LL-37 itself [1]. Imbalances in the expression levels of this peptide could lead to inflammatory diseases such as Crohn’s disease [5,6] or skin lesions driven by epithelial tissue inflammation [7]; and being an HDP, LL-37 has antimicrobial activity and diverse immunomodulatory effects [8,9]. Hence, diverse efforts have been made to identify small peptides derived from LL-37 with the same activities [8–11] and/or understand the importance of specific amino acid residues in their sequence [12–14].

Albeit the clinical and biotechnological importance of LL-37, the effects of missense single nucleotide polymorphisms (SNPs) on the LL-37 are unknown. Testing these mutations *in vitro* could be expensive, and nowadays, computational analysis have become an alternative [15]. Initially, the missense SNPs are subjected to predictions for classifying them as deleterious or neutral by a number of prediction tools. However, these tools have presented limitations in benchmarking analyses using specific proteins [16–19] and for this reason, further evaluation by molecular dynamics simulations is required. This approach has been applied by our group to study SNP effects in different peptides, including guanylins [20–22] and defensins [23,24]; and also larger proteins, including growth hormone receptor [25], apolipoprotein E [26] and glucocorticoid receptor [27]. Besides, other groups applied molecular dynamics to assess the effects of mutations on aurora-A kinase [28], lamin A/C protein [29] and PncA protein [30]. In the present work, we applied a similar strategy to assess the effects of missense SNPs on LL-37 structure, showing the SNPs did not affect the residues from the active core of LL-37; however they could influence the antimicrobial activity by altering the positive net charge, helix dipole and/or the α-helix structure.

## 2 Results

### 2.1 Selected SNPs alter only the terminal portions of LL-37

The *CAMP* gene encodes a protein sequence containing 170 amino acid residues. The last 37 residues correspond to the mature LL-37, and therefore, the selected SNPs were restricted to this last portion (Figure 1A). A total of 11 validated SNPs occurring within the mature sequence were identified. Once this is a feasible number of SNPs to analyze, no additional filters were applied for selection. On the N-terminal portion there are six modifications, while on C-terminal portion presented the remaining five (Figure 1B). The central portion (^13^IGKEFKRIVQRIKDFL^28^) was not affected. In addition, the amphipathic distribution is not affected, because the positions with mutations either they are near to the terminals (Figure 1B) or they occur on the borders of hydrophobic/hydrophilic faces (Figure 1C).

**Figure 1.**
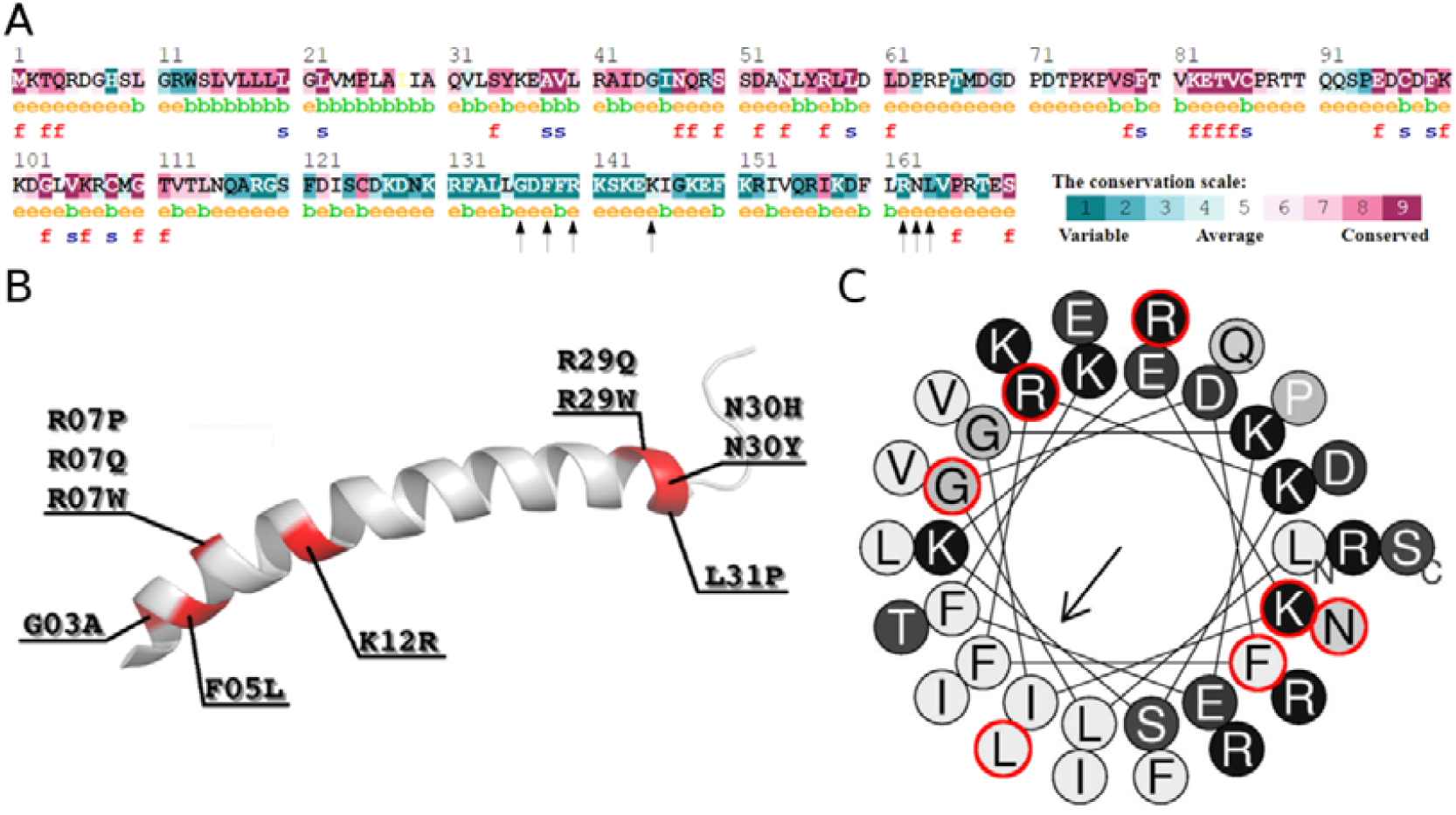
LL-37 point mutations overview. (A) Consurf amino acid conservation pattern. Consurf server searches for homologous sequences to the query sequence and build patterns based on amino acid conservation. The patterns are shown by a color scale, in which the grade of 1 (turquoise) represent the residues with high variability, and the grade of 9 (magenta) represent the residues that are highly conserved among homologous. The variability of an amino acid is usually associated with fundamental structural and/or functional activity. These characteristics were predicted and represented by the letters f- a predicted functional residue (highly conserved and exposed) and s- a predicted structural residue (highly conserved and buried). The letters e and b represent residues exposed or buried, respectively, according to the neural network algorithm. The arrows indicate the changed residues. All LL-37 point mutations are in low conserved regions. (B) LL-37 three-dimensional structure. This peptide presents a slightly curved α-helix from residues Leu^1^-Leu^31^. The point mutation positions caused by SNPs are highlighted in red. (C) Helical wheel projection of LL-37. The residues are colored in shades of gray from the more hydrophobic (light grey) to more hydrophilic (dark grey). The mutated residues are highlighted in by a red circle. The arrow indicates the length and the direction of the hydrophobic moment vector.

### 2.2 Low amino acid conservation hinders a proper SNP effect prediction

The cathelin domain is more conserved than the mature LL-37, which presents a low conservation degree; and therefore, all selected mutations occurred on low conserved positions (Figure 1A). The absence of conservation is reflected on SNP effect predictions, where virtually all mutations were classified as neutral by different algorithms (Table 1).

**Table 1.**
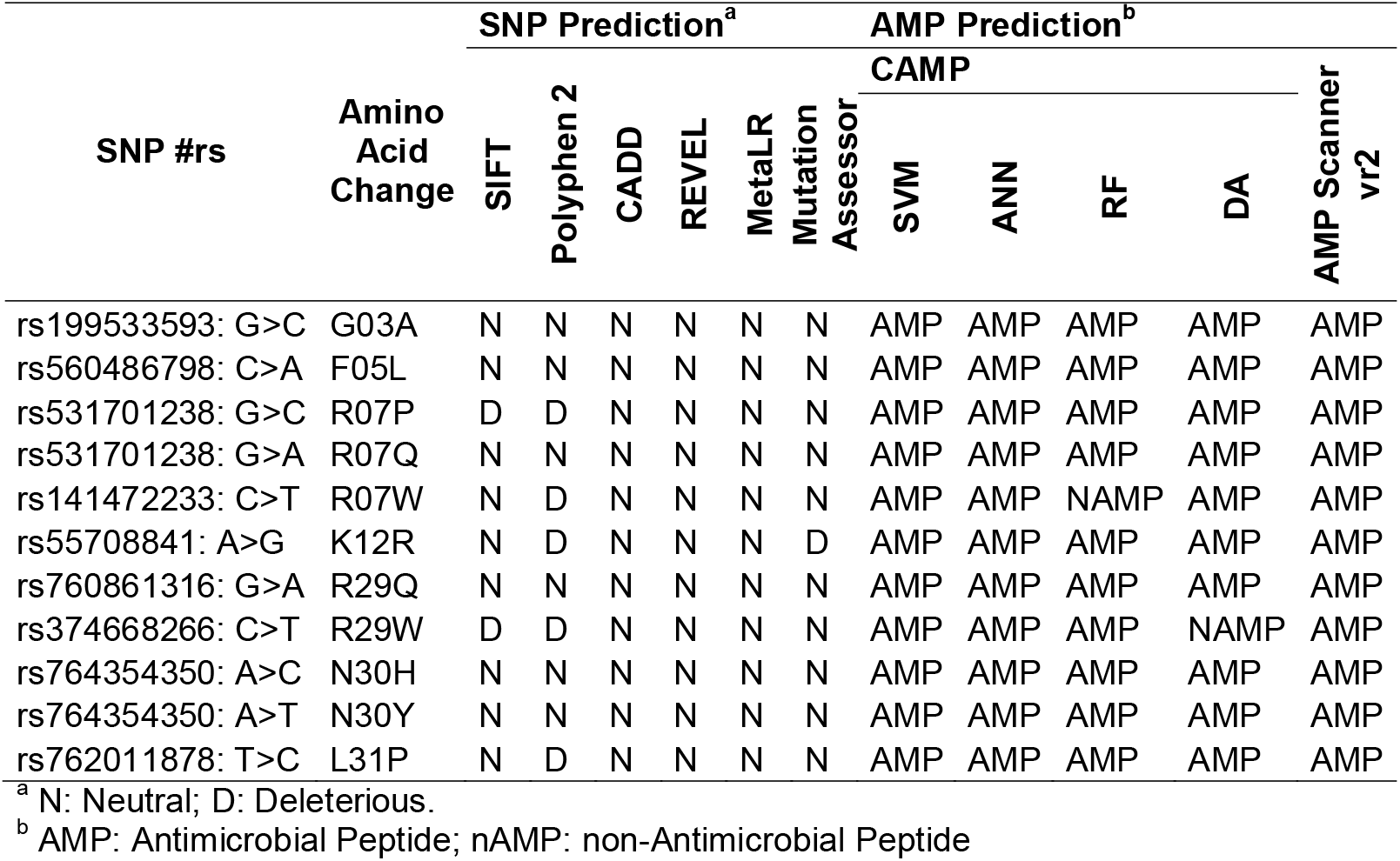
Prediction analysis of SNP effects and antimicrobial activity.

### 2.3 Antimicrobial activity is predicted to be maintained for all variants

Because antimicrobial activity is one of the main LL-37 functions, the selected variants were subjected to antimicrobial activity predictors, where virtually all variants were classified as AMP in all algorithms, except for R07W and R29W, which had negative predictions according to CAMP Random Forest and CAMP Discriminant Analysis predictions, respectively (Table 1). Considering the variants are 97% identical to the wild-type peptide and the predictors are binary classifiers, these prediction results are not surprising. However, these point mutations generate modifications on the physicochemical properties (Table 2), in particular on charged residues, which in turn could affect the antimicrobial activity. The R07P, R07Q, R07W, R29Q and R29W mutations had a charge reduction; while N30H mutation had a charge increase (Table 2). In addition, the majority of mutations induced a slight increase in the hydrophobicity and a slight reduction in the hydrophobic moment (Table 2).

**Table 2.** Physicochemical properties of LL-37 variants.

| Variant | Hydrophobicity | Hydrophobic Moment | Net Charge |
| --- | --- | --- | --- |
| Wild-Type | -0.347 | 0.361 | 6 |
| G03A | -0.345 | 0.362 | 6 |
| F05L | -0.349 | 0.361 | 6 |
| R07P | -0.301 | 0.355 | 5 |
| R07Q | -0.317 | 0.357 | 5 |
| R07W | -0.289 | 0.355 | 5 |
| K12R | -0.366 | 0.371 | 6 |
| R29Q | -0.317 | 0.338 | 5 |
| R29W | -0.289 | 0.318 | 5 |
| N30H | -0.341 | 0.358 | 7 |
| N30Y | -0.329 | 0.353 | 6 |
| L31P | -0.364 | 0.346 | 6 |

### 2.4 R07Q and N30Y variants changes the LL-37 structure

Despite both kind of predictions based on the primary sequence (Table 1) indicated the variations cause no changes on the LL-37 activity, we applied molecular dynamics simulations to perform a more refined assessment, regarding the structure-activity relationship. Because the models were constructed using the LL-37 structure as template, their structures were composed by a curved α-helix comprising the residues Leu^1^ to Leu^31^, followed by a short tail, virtually identical to that displayed on Figure 1B. All models were considered acceptable, with a slight variation on the assessments (Table 3).

**Table 3.** Summary of three-dimensional models validation.

| Variant | DOPE | Z-Score<br>(Prosa II) | Ramachandran Plot (%) |  | QMEAN4 |
| --- | --- | --- | --- | --- | --- |
|  |  |  | Favoured | Allowed |  |
| G03A | -2874.147 | -1.45 | 97 | 3 | 0.97 |
| F05L | -2843.359 | -1.35 | 93.8 | 3.1 | 0.4 |
| R07P | -2723.808 | -0.87 | 96.8 | 3.2 | 0.34 |
| R07Q | -2823.960 | -1.42 | 100 | 0 | 0.92 |
| R07W | -2890.820 | -0.98 | 96.9 | 3.1 | 0.95 |
| K12R | -2831.043 | -1.49 | 96.9 | 3.1 | 0.94 |
| R29Q | -2815.620 | -1.31 | 93.8 | 3.1 | 0.16 |
| R29W | -2950.267 | -1.12 | 96.9 | 3.1 | 1.08 |
| N30H | -2844.895 | -1.52 | 96.9 | 0 | 0.44 |
| N30Y | -2886.529 | -1.34 | 100 | 0 | 0.65 |
| L31P | -2742.228 | -0.76 | 96.8 | 0 | 0.73 |

On molecular dynamics simulation on saline solution, we could observe a partial unfolding process, where the root mean square deviations reached values between 4 and 5 Å (Figure 2A). This deviation could be a result of C-Terminal loop moves along the simulation, as the deviation from ideal helix was kept below 4 Å (Figure 2B). However, the R07Q and N30Y variants did not kept the initial α-helix, which could be observed from the high values from RMSD and also the deviation from ideal helix. In fact, these variants populated a different cluster on essential dynamics (Figure 2C). Despite the R07W variant showed a peak of deviation from ideal helix (Figure 2B), this variant populated the same cluster from wild-type structure (data not shown).

**Figure 2.**
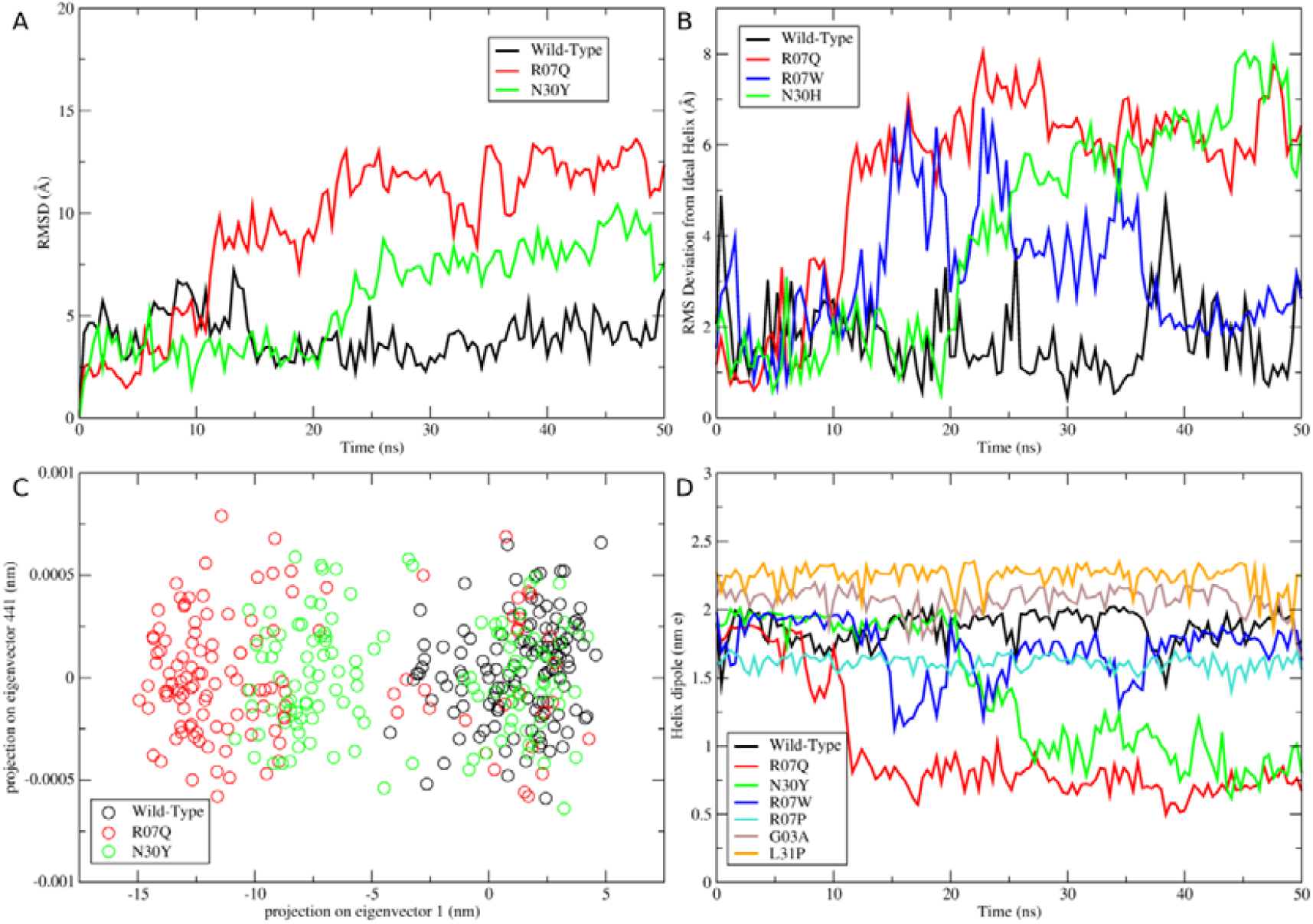
Summary of molecular dynamics results. In all panels, variants with similar behavior of wild-type LL-37 were omitted for clarity. (A) Backbone’s RMSD evolution along the simulation. The wild-type LL-37 maintained an RMSD value between 4 and 5 Å, keeping the linear α-helix structure; while the R07Q and N30Y variants did not kept the linear α-helix structure, presenting a kink α-helix structure. (B) RMS Deviation from ideal α-helix. The R07Q and N30Y variants clearly deviated from an ideal α-helix structure, while the R07W variant refolded into an α-helix after a unfolding process (C) Projection of the motion of LL-37 in phase space along the 1^st^ and the 441^st^ eigenvectors. Because the LL-37 three-dimensional structure is its own secondary structure, only the 1^st^ eigenvector showed some variation, the remaining were close to zero. From this projection two clusters could be observed, the first one in around of coordinates [0,0], where the wild-type and other variants fell, and the second one in around of coordinates [-10, 0], where R07Q and N30Y variants fell, representing a different structure. (D) Helix dipole along the simulations. The helix dipole is important to peptide-membrane interactions; four variants presented a reduction in the helix dipole (R07P, R07Q, R07W and N30Y variants), while the variants G03A and L31P increased this property.

### 2.5 G03A, R07P, R07Q, R07W, N30Y and L31P mutations alter the helix dipole

The α-helix structure is a key to LL-37 activity and in this context, the helix dipole is important for membrane interactions. Because R07Q and N30Y adopt different structures from the wild-type and other variants, their helix dipole is reduced by a half (Figure 2D). Interestingly, we could observe alterations on such properties even in variants that kept a similar structure of wild-type, where G03A and L31P increased their helix dipoles, while R07P and R07W showed a reduction on such property (Figure 2D). Therefore, the G03A and L31P could have a stronger interaction with membranes due to the increased helix dipole, while R07Q and N30Y, could have such interaction compromised, due to the helix disruption and consequent reduction of helix dipole.

## 3 Discussion

Due to its multiple functions and roles, the cathelicidin LL-37 has been the subject of a number of studies in the last years. On the one hand, its antimicrobial activity has led to development of short AMPs derived from its active core [8–11] and artificial mutations have led to the understanding of the importance of each amino acid residue for the antimicrobial activity [12–14]; and on the other hand, the LL-37 role in immune system has led to the discovery of its relationship with some diseases [5–7]. In such context, the present work brings the opportunity to study the structure-activity relationships together with the genetic background under the light of antimicrobial activity.

From an evolutionary perspective, producing antibiotic molecules is imperative for survival of an organism [1,2]. In humans LL-37 is not the unique, but is one of the most remarkable antimicrobial molecules. Therefore, the presence of mutations that abolish this activity would be counterintuitive, which lead us to the result that no SNPs affecting the central residues of LL-37 were identified. This finding is in alignment with previous studies that had indicated KR12, the fragment containing residues 18-29, as the minimal active fragment [10,11]; and in fact, only the R29Q and R29W affect this portion (Figure 1A and B).

Considering the active core is preserved in the majority of the variants and the hydrophobic/hydrophilic faces are kept (Figure 1C), with slight hydrophobic moment variations (Table 2), the antimicrobial activity maintenance is expected. Indeed, the variants were predicted as antimicrobial (Table 1). Conversely, when the net charge is reduced, even in larger LL-37 fragments, the potency of antimicrobial activity is reduced [12–14]. Hence, the mutations involving Arg residues (R07P, R07Q, R07W, R29Q and R29W) should have their antimicrobial potency reduced, because these variants have a net charge of +5, while the wild-type has +6 (Table 2).

According to this reasoning, these mutations are probably deleterious, in contrast to the SNP effect predictions, which indicated the variants are neutral (Table 1). Albeit contrasting, the SNP effect prediction is biased due to the conservation pattern of LL-37 precursor (Figure 1A). An accurate SNP effect prediction is hindered because those programs extract some kind of information from multiple sequence alignments among homologous sequences [31]. In addition, disease-causing missense SNPs tend to occur at evolutionarily conserved positions that have an essential role in the structure and/or function of the encoded protein [32]. Therefore such kind of pattern could not be extracted for mature LL-37 (Figure 1A).

In the last years, the validity of applying such predictors has been put in check after benchmarking analysis for both antimicrobial activity [33] and SNP effect [16–19] predictions. Nevertheless, this kind of prediction has been used as a filter to select variants for more refined analysis [20,21,26]. In our previous work with other human AMPs, we observed a similar pattern of neutral predictions, however, the structural analysis revealed the deleterious character of some variants [24]. Hence, we simulated all selected variants, despite the neutral predictions. The R07Q and N30Y variants clearly changed the LL-37 structure (Figure 2A-C), which could affect their antimicrobial activities, turning them potential deleterious mutations. Once the structure was altered, these mutations have their helix dipole reduced (Figure 2D), which could hinder their membrane interactions. The other mutations in Arg^7^ also reduced the helix dipole (Figure 2D), despite they kept the α-helix, thus they can also have a deleterious effect. Finally, two mutations (G03A and L31P) increased the helix dipole, which in turn could improve their membrane interactions [2,34], turning them toxic peptides and therefore deleterious mutations. Previously, we applied solvation potential energy analysis to assess the effects of point mutations on human defensins (HD5 and HBD1) [23,24]; nevertheless the same analysis for LL-37 variants was inconclusive (data not shown), which could be related to differences in their mechanism of actions, because LL-37 and defensins belong to different families of AMPs.

## 4 Conclusion

LL-37 antimicrobial activity is very sensitive to point mutations [12–14]. And due to its multiple activities in human body, point mutations on the active core of this peptide may lead loss of function [12–14] and therefore removed from population by negative selection. From the point mutations induced by SNPs, none of them affect the LL-37 core, but they could lead to changes in antimicrobial activity. From these mutations, only F05L and K12R seem to be neutral. The R07P, R07W, R29Q, R29W variants could have their potency reduced due to the reduced charge, while the G03A, L31P and N30H variants could have an increased activity, the first two due to the increase on the helix dipole and the last one due to an increase in the net charge. Nevertheless, the most remarkable mutations were R07Q and N30Y, which presented structural modifications, and in particular R07Q, which also loses one positive charge. The *in vitro* and *in vivo* assessments of such variants is essential to generate an in depth knowledge of the functional and structural effects of these SNPs and to perform further genetic association studies. Our results indicated that despite the mutations did not alter the residues from LL-37 active core, they could influence the antimicrobial activity and consequently, could be involved in inflammatory diseases.

## 5 Material and Methods

### 5.1 Databases and SNPs selection

The SNPs were retrieved from dbSNP database (build 153) using the Variation Viewer navigator from the NCBI (https://www.ncbi.nlm.nih.gov/variation/view/) [35]. The missense variant filter was applied and only the *CAMP* SNPs located in the mature sequence, and those validated by at least one of the following criteria were selected: SNPs obtained through targeted deep (50-100X) exome sequencing; a combination of this and low coverage (2-6X) whole-genome sequencing; SNPs with multiple, independent submissions to the refSNP cluster; SNPs validated by frequency or genotype data; SNPs genotyped by HapMap; or SNPs with frequency on 1000 Genomes Project. The LL-37 sequence and the protein structure files of the mature peptide (PDB ID: 2K6O)[11] were obtained from the RCSB Protein Data Bank. The precursor sequence was obtained from NCBI (P49913.1).

### 5.2 SNP effect and antimicrobial activity predictions

The mature LL-37 sequence and their variants were subjected to antimicrobial activity prediction using the deep learning based predictor AMP Scanner vr2 [36] and four algorithms from Collection of Antimicrobial Peptides (CAMP), including support vector machine, artificial neural network, discriminant analysis and random forest [37]. The SNP effect predictions were retrieved from Human genome (GRCh37), from Ensembl, including SIFT [38], PolyPhen 2 [39], Mutation Assessor [40], CADD [41], REVEL [42] and MetaLR [43].

### 5.3 Physicochemical properties calculation

Physicochemical properties, including net charge, average hydrophobicity and hydrophobic moment were calculated using a PERL script^1^. For net charge calculation, histidine residues were considered as positively charged; the average hydrophobicity and hydrophobic moment were calculated using the Eisenberg Scale [44] and the average hydrophobic moment was calculated using Eisenberg’s equation [44]:

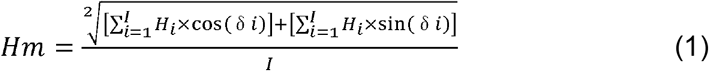

Where δ represents the angle between the amino acid side chains (100° for α-helix, on average); *i* represents the residue number in the position *i* from the sequence; H_*i*_ represents the *i*^th^ amino acid’s hydrophobicity on a hydrophobicity scale; and *I* represents the total number of residues. Heliquest server [45] was used to draw the helical wheel projection.

### 5.4 Evolutionary conservation analysis

The ConSurf [46] is a webserver that based on the phylogenetic relations between homologous sequences. This tool can estimate the evolutionary conservation of amino acid positions in a protein molecule. Using the precursor sequence obtained from the National Center of Biotechnology Information (P49913.1), ConSurf, in ConSeq mode, carried out a search for close homologous sequences using CSI-BLAST (3 iterations and 0.0001 e-value cutoff) against the UNIREF-90 protein database. The settings were set as follows, maximum number of homologs as 150 and a 35 minimal to 95 maximal percentage ID between sequences. Multiple sequence alignment and calculation methods were used as default (MAFFT-L-INS-I and Bayesian). The sequences were then clustered and highly similar sequences removed using CD-HIT [47]. The empirical Bayesian algorithm was used to compute the position-specific conservation scores.

### 5.5 Molecular modelling

One hundred molecular models for each variant were constructed by comparative molecular modeling by means of MODELLER 9.17 [48], using the LL-37 structure (PDB ID: 2K6O) [11]. The models were constructed using the default methods of automodel and environ classes from MODELLER. The final models were selected according to the discrete optimized protein energy score (DOPE score). This score assesses the energy of the models and indicates the best probable structures. The best models were evaluated by means of QMEAN [49], PROSA II [50] and PROCHECK [51]. PROCHECK checks the stereochemical quality of a protein structure by means of the Ramachandran plot, where good quality models are expected to have more than 90% of amino acid residues in most favored and additional allowed regions, while PROSA II indicates the fold quality; and QMEAN, in turn, uses a scoring function that combines six structural descriptors to estimate the absolute quality of a protein structure. Structural visualization was done in PyMOL (http://www.pymol.org).

### 5.6 Molecular dynamics simulation

The molecular dynamics simulation of the wild-type and variants were performed in the GROMACS 4.6 package [52] using the CHARMM force field [53]. The simulations were performed in a saline solution (0.2 M NaCl) where the structures were immersed in water cubic boxes with a 0.8 nm distance between the protein and the edge of the box. Geometry of water molecules was constrained by using the SETTLE algorithm [54]. All atom bond lengths were linked by using the LINCS algorithm [55]. Electrostatic corrections were made by Particle Mesh Ewald algorithm [56], with a threshold of 1.4 nm to minimize the computational time. The same cut-off radius was also used for van der Waals interactions. The steepest descent algorithm was applied to minimize system energy for 50,000 steps. After that, the system temperature was normalized to 310 K for 100 ps, using the velocity rescaling thermo-stat (NVT ensemble). Next, the system pressure was normalized to 1 bar for 100 ps, using the Parrinello–Rahman barostat (NPT ensemble). The system with temperature normalized to 310 K and pressure normalized to 1 bar were simulated for 50 ns.

### 5.7 Analyses of molecular dynamics trajectories

Molecular dynamics simulations were analyzed by means of the backbone root mean square deviation (RMSD), RMS deviation form ideal helix and helix dipole using the g_rms, g_helix built in functions of the GROMACS package. The RMSD that used the GROMACS group 4 (backbone) to analyses, the functions used the GROMACS group 1 (protein). The principal component analysis (PCA) was used to analyze and visualize the protein motion in the simulation. For that, the protein movement is extracted and split in different components. For visualization, the 1^st^ and the 441^st^ eigenvectors were plot against each other. The PCA, or essential dynamics analysis, were done using the g_covar and g_anaeig functions.

## 6 Acknowledgements

This work was supported by CNPq (Conselho Nacional de Desenvolvimento Científico e Tecnológico); CAPES (Coordenação de Aperfeiçoamento de Pessoal de Nível Superior) and FAPDF (Fundação de Amparo a Pesquisa do Distrito Federal).

## Footnotes

1 Available at https://github.com/williamfp7/AMPProperties

## Notes

https://github.com/williamfp7/AMPProperties

### Competing Interest Statement

The authors have declared no competing interest.

### Summary of Updates

Revised version after peer-review process.

## References

[1] O.N. Silva, W.F. Porto, S.M. Ribeiro, I. Batista, O.L. Franco, Host-defense peptides and their potential use as biomarkers in human diseases, Drug Discov. Today. 23 (2018) 1666–1671. https://doi.org/10.1016/j.drudis.2018.05.024.

[2] N. Sitaram, R. Nagaraj, Interaction of antimicrobial peptides with biological and model membranes: Structural and charge requirements for activity, Biochim. Biophys. Acta - Biomembr. 1462 (1999) 29–54. https://doi.org/10.1016/S0005-2736(99)00199-6.

[3] M. Zanetti, R. Gennaro, D. Romeo, Cathelicidins: a novel protein family with a common proregion and a variable C-terminal antimicrobial domain, FEBS Lett. 374 (1995) 1–5. https://doi.org/10.1016/0014-5793(95)01050-O.

[4] E.M. Kościuczuk, P. Lisowski, J. Jarczak, N. Strzałkowska, A. Jóźwik, J. Horbańczuk, J. Krzyżewski, L. Zwierzchowski, E. Bagnicka, Cathelicidins: family of antimicrobial peptides. A review, Mol. Biol. Rep. 39 (2012) 10957–10970. https://doi.org/10.1007/s11033-012-1997-x.

[5] A.F. Gombart, T. Saito, H.P. Koeffler, Exaptation of an ancient Alu short interspersed element provides a highly conserved vitamin D-mediated innate immune response in humans and primates., BMC Genomics. 10 (2009) 321. https://doi.org/10.1186/1471-2164-10-321.

[6] M.B. Lowry, C. Guo, N. Borregaard, A.F. Gombart, Regulation of the human cathelicidin antimicrobial peptide gene by 1α,25-dihydroxyvitamin D3 in primary immune cells., J. Steroid Biochem. Mol. Biol. 143 (2014) 183–91. https://doi.org/10.1016/j.jsbmb.2014.02.004.

[7] T.-T. Wang, L.E. Tavera-Mendoza, D. Laperriere, E. Libby, N.B. MacLeod, Y. Nagai, V. Bourdeau, A. Konstorum, B. Lallemant, R. Zhang, S. Mader, J.H. White, Large-scale in silico and microarray-based identification of direct 1,25-dihydroxyvitamin D3 target genes., Mol. Endocrinol. 19 (2005) 2685–95. https://doi.org/10.1210/me.2005-0106.

[8] T. Sigurdardottir, P. Andersson, M. Davoudi, M. Malmsten, A. Schmidtchen, M. Bodelsson, In silico identification and biological evaluation of antimicrobial peptides based on human cathelicidin LL-37., Antimicrob. Agents Chemother. 50 (2006) 2983–9. https://doi.org/10.1128/AAC.01583-05.

[9] G. Wang, M. Elliott, A.L. Cogen, E.L. Ezell, R.L. Gallo, R.E.W. Hancock, Structure, dynamics, and antimicrobial and immune modulatory activities of human LL-23 and its single-residue variants mutated on the basis of homologous primate cathelicidins., Biochemistry. 51 (2012) 653–64. https://doi.org/10.1021/bi2016266.

[10] X. Li, Y. Li, H. Han, D.W. Miller, G. Wang, Solution structures of human LL-37 fragments and NMR-based identification of a minimal membrane-targeting antimicrobial and anticancer region., J. Am. Chem. Soc. 128 (2006) 5776–85. https://doi.org/10.1021/ja0584875.

[11] G. Wang, Structures of human host defense cathelicidin LL-37 and its smallest antimicrobial peptide KR-12 in lipid micelles., J. Biol. Chem. 283 (2008) 32637–43. https://doi.org/10.1074/jbc.M805533200.

[12] S. Gunasekera, T. Muhammad, A.A. Strömstedt, K.J. Rosengren, U. Göransson, Alanine and Lysine Scans of the LL-37-Derived Peptide Fragment KR-12 Reveal Key Residues for Antimicrobial Activity, ChemBioChem. 19 (2018) 931–939. https://doi.org/10.1002/cbic.201700599.

[13] G. Wang, R.F. Epand, B. Mishra, T. Lushnikova, V.C. Thomas, K.W. Bayles, R.M. Epand, Decoding the Functional Roles of Cationic Side Chains of the Major Antimicrobial Region of Human Cathelicidin LL-37, Antimicrob. Agents Chemother. 56 (2012) 845–856. https://doi.org/10.1128/AAC.05637-11.

[14] B. Mishra, R.F. Epand, R.M. Epand, G. Wang, Structural location determines functional roles of the basic amino acids of KR-12, the smallest antimicrobial peptide from human cathelicidin LL-37, RSC Adv. 3 (2013) 19560. https://doi.org/10.1039/c3ra42599a.

[15] H. Shen, JieLJ; Deininger Prescott.; Zhao, Applications of computational algorithm tools to identify functional SNPs in cytokine genes, Cytokine. 35 (2006) 62–66.

[16] K.C.D.V. Ferreira, L.F. Fialho, O.L. Franco, S.A. de Alencar, W.F. Porto, Benchmarking analysis of deleterious SNP prediction tools on CYP2D6 enzyme, Chem. Biol. Drug Des. 96 (2020). https://doi.org/10.1111/cbdd.13676.

[17] D.G. Grimm, C.-A. Azencott, F. Aicheler, U. Gieraths, D.G. MacArthur, K.E. Samocha, D.N. Cooper, P.D. Stenson, M.J. Daly, J.W. Smoller, L.E. Duncan, K.M. Borgwardt, The evaluation of tools used to predict the impact of missense variants is hindered by two types of circularity., Hum. Mutat. 36 (2015) 513–23. https://doi.org/10.1002/humu.22768.

[18] S. Hicks, D.A. Wheeler, S.E. Plon, M. Kimmel, Prediction of missense mutation functionality depends on both the algorithm and sequence alignment employed, Hum. Mutat. (2011). https://doi.org/10.1002/humu.21490.

[19] C. Rodrigues, A. Santos-Silva, E. Costa, E. Bronze-da-Rocha, Performance of In Silico Tools for the Evaluation of UGT1A1 Missense Variants., Hum. Mutat. 36 (2015) 1215–25. https://doi.org/10.1002/humu.22903.

[20] W.F. Porto, O.L. Franco, S.A. Alencar, Computational analyses and prediction of guanylin deleterious SNPs, Peptides. 69 (2015) 92–102. https://doi.org/10.1016/j.peptides.2015.04.013.

[21] A.C.S. Marcolino, W.F. Porto, Á.S. Pires, O.L. Franco, S.A. Alencar, Structural impact analysis of missense SNPs present in the uroguanylin gene by long-term molecular dynamics simulations., J. Theor. Biol. 410 (2016) 9–17. https://doi.org/10.1016/j.jtbi.2016.09.008.

[22] Á.S. Pires, W.F. Porto, P.O. Castro, O.L. Franco, S.A. Alencar, Theoretical structural characterization of lymphoguanylin: A potential candidate for the development of drugs to treat gastrointestinal disorders., J. Theor. Biol. 419 (2017) 193–200. https://doi.org/10.1016/j.jtbi.2017.02.016.

[23] W.F. Porto, D.O. Nolasco, Á.S. Pires, G.R. Fernandes, O.L. Franco, S.A. Alencar, HD5 and HBD1 variants’ solvation potential energy correlates with their antibacterial activity against Escherichia coli, Biopolymers. 106 (2016) 43–50. https://doi.org/10.1002/bip.22763.

[24] W.F. Porto, D.O. Nolasco, Á.S. Pires, R.W. Pereira, O.L. Franco, S.A. Alencar, Prediction of the impact of coding missense and nonsense single nucleotide polymorphisms on HD5 and HBD1 antibacterial activity against Escherichia coli, Biopolymers. 106 (2016) 633–644. https://doi.org/10.1002/bip.22866.

[25] W.F. Porto, F.A. Marques, H.B. Pogue, M.T. de Oliveira Cardoso, M.G.R. do Vale, Á. da Silva Pires, O.L. Franco, S.A. de Alencar, R. Pogue, Computational Investigation of Growth Hormone Receptor Trp169Arg Heterozygous Mutation in a Child With Short Stature, J. Cell. Biochem. 118 (2017) 4762–4771. https://doi.org/10.1002/jcb.26144.

[26] A.S. Pires, W.F. Porto, O.L. Franco, S.A. Alencar, In silico analyses of deleterious missense SNPs of human apolipoprotein E3., Sci. Rep. 7 (2017) 2509. https://doi.org/10.1038/s41598-017-01737-w.

[27] L.L.S. Monteiro, O.L. Franco, S.A. Alencar, W.F. Porto, Deciphering the structural basis for glucocorticoid resistance caused by missense mutations in the ligand binding domain of glucocorticoid receptor, J. Mol. Graph. Model. 92 (2019) 216–226. https://doi.org/10.1016/j.jmgm.2019.07.020.

[28] A. Kumar, R. Purohit, Use of Long Term Molecular Dynamics Simulation in Predicting Cancer Associated SNPs, PLoS Comput. Biol. 10 (2014) e1003318. https://doi.org/10.1371/journal.pcbi.1003318.

[29] V. Rajendran, R. Purohit, R. Sethumadhavan, In silico investigation of molecular mechanism of laminopathy caused by a point mutation (R482W) in lamin A/C protein., Amino Acids. 43 (2012) 603–15. https://doi.org/10.1007/s00726-011-1108-7.

[30] V. Rajendran, R. Sethumadhavan, Drug resistance mechanism of PncA in Mycobacterium tuberculosis., J. Biomol. Struct. Dyn. 32 (2014) 209–21. https://doi.org/10.1080/07391102.2012.759885.

[31] P.C. Ng, S. Henikoff, Predicting the effects of amino acid substitutions on protein function., Annu. Rev. Genomics Hum. Genet. 7 (2006) 61–80. https://doi.org/10.1146/annurev.genom.7.080505.115630.

[32] M.P. Miller, S. Kumar, Understanding human disease mutations through the use of interspecific genetic variation., Hum. Mol. Genet. 10 (2001) 2319–28.

[33] W.F. Porto, Á.S. Pires, O.L. Franco, Antimicrobial activity predictors benchmarking analysis using shuffled and designed synthetic peptides., J. Theor. Biol. 426 (2017) 96–103. https://doi.org/10.1016/j.jtbi.2017.05.011.

[34] P. Juvvadi, S. Vunnam, R.B. Merrifield, Synthetic Melittin, Its Enantio, Retro, and Retroenantio Isomers, and Selected Chimeric Analogs: Their Antibacterial, Hemolytic, and Lipid Bilayer Action, J. Am. Chem. Soc. 118 (1996) 8989–8997. https://doi.org/10.1021/ja9542911.

[35] S.T. Sherry, M.H. Ward, M. Kholodov, J. Baker, L. Phan, E.M. Smigielski, K. Sirotkin, dbSNP: the NCBI database of genetic variation., Nucleic Acids Res. 29 (2001) 308–11.

[36] D. Veltri, U. Kamath, A. Shehu, Deep learning improves antimicrobial peptide recognition, Bioinformatics. 34 (2018) 2740–2747. https://doi.org/10.1093/bioinformatics/bty179.

[37] S. Thomas, S. Karnik, R.S. Barai, V.K. Jayaraman, S. Idicula-Thomas, CAMP: a useful resource for research on antimicrobial peptides., Nucleic Acids Res. 38 (2010) D774–80. https://doi.org/10.1093/nar/gkp1021.

[38] P. Kumar, S. Henikoff, P.C. Ng, Predicting the effects of coding non-synonymous variants on protein function using the SIFT algorithm., Nat. Protoc. 4 (2009) 1073–81. https://doi.org/10.1038/nprot.2009.86.

[39] I.A. Adzhubei, S. Schmidt, L. Peshkin, V.E. Ramensky, A. Gerasimova, P. Bork, A.S. Kondrashov, S.R. Sunyaev, A method and server for predicting damaging missense mutations., Nat. Methods. 7 (2010) 248–9. https://doi.org/10.1038/nmeth0410-248.

[40] B. Reva, Y. Antipin, C. Sander, Predicting the functional impact of protein mutations: application to cancer genomics., Nucleic Acids Res. 39 (2011) e118. https://doi.org/10.1093/nar/gkr407.

[41] P. Rentzsch, D. Witten, G.M. Cooper, J. Shendure, M. Kircher, CADD: predicting the deleteriousness of variants throughout the human genome, Nucleic Acids Res. 47 (2019) D886–D894. https://doi.org/10.1093/nar/gky1016.

[42] N.M. Ioannidis, J.H. Rothstein, V. Pejaver, S. Middha, S.K. McDonnell, S. Baheti, A. Musolf, Q. Li, E. Holzinger, D. Karyadi, L.A. Cannon-Albright, C.C. Teerlink, J.L. Stanford, W.B. Isaacs, J. Xu, K.A. Cooney, E.M. Lange, J. Schleutker, J.D. Carpten, I.J. Powell, O. Cussenot, G. Cancel-Tassin, G.G. Giles, R.J. MacInnis, C. Maier, C.-L. Hsieh, F. Wiklund, W.J. Catalona, W.D. Foulkes, D. Mandal, R.A. Eeles, Z. Kote-Jarai, C.D. Bustamante, D.J. Schaid, T. Hastie, E.A. Ostrander, J.E. Bailey-Wilson, P. Radivojac, S.N. Thibodeau, A.S. Whittemore, W. Sieh, REVEL: An Ensemble Method for Predicting the Pathogenicity of Rare Missense Variants, Am. J. Hum. Genet. 99 (2016) 877–885. https://doi.org/10.1016/j.ajhg.2016.08.016.

[43] C. Dong, P. Wei, X. Jian, R. Gibbs, E. Boerwinkle, K. Wang, X. Liu, Comparison and integration of deleteriousness prediction methods for nonsynonymous SNVs in whole exome sequencing studies., Hum. Mol. Genet. 24 (2015) 2125–37. https://doi.org/10.1093/hmg/ddu733.

[44] D. Eisenberg, R.M. Weiss, T.C. Terwilliger, W. Wilcox, Hydrophobic moments and protein structure, Faraday Symp. Chem. Soc. 17 (1982) 109. https://doi.org/10.1039/fs9821700109.

[45] R. Gautier, D. Douguet, B. Antonny, G. Drin, HELIQUEST: a web server to screen sequences with specific alpha-helical properties., Bioinformatics. 24 (2008) 2101–2. https://doi.org/10.1093/bioinformatics/btn392.

[46] G. Celniker, G. Nimrod, H. Ashkenazy, F. Glaser, E. Martz, I. Mayrose, T. Pupko, N. Ben-Tal, ConSurf: Using Evolutionary Data to Raise Testable Hypotheses about Protein Function, Isr. J. Chem. 53 (2013) 199–206. https://doi.org/10.1002/ijch.201200096.

[47] W. Li, A. Godzik, Cd-hit: a fast program for clustering and comparing large sets of protein or nucleotide sequences., Bioinformatics. 22 (2006) 1658–9. https://doi.org/10.1093/bioinformatics/btl158.

[48] N. Eswar, B. Webb, M.A. Marti-Renom, M.S. Madhusudhan, D. Eramian, M.-Y. Shen, U. Pieper, A. Sali, Comparative protein structure modeling using Modeller., Curr. Protoc. Bioinforma. Chapter 5 (2006) Unit-5.6. https://doi.org/10.1002/0471250953.bi0506s15.

[49] P. Benkert, M. Biasini, T. Schwede, Toward the estimation of the absolute quality of individual protein structure models, Bioinformatics. 27 (2011) 343–350. https://doi.org/10.1093/bioinformatics/btq662.

[50] M. Wiederstein, M.J. Sippl, ProSA-web: interactive web service for the recognition of errors in three-dimensional structures of proteins., Nucleic Acids Res. 35 (2007) W407–10. https://doi.org/10.1093/nar/gkm290.

[51] R.A. Laskowski, M.W. MacArthur, D.S. Moss, J.M. Thornton, PROCHECK: a program to check the stereochemical quality of protein structures, J. Appl. Crystallogr. 26 (1993) 283–291. https://doi.org/10.1107/S0021889892009944.

[52] B. Hess, C. Kutzner, D. van der Spoel, E. Lindahl, GROMACS 4: Algorithms for Highly Efficient, Load-Balanced, and Scalable Molecular Simulation, J. Chem. Theory Comput. 4 (2008) 435–447. https://doi.org/10.1021/ct700301q.

[53] K. Vanommeslaeghe, E. Hatcher, C. Acharya, S. Kundu, S. Zhong, J. Shim, E. Darian, O. Guvench, P. Lopes, I. Vorobyov, A.D. Mackerell, CHARMM general force field: A force field for drug-like molecules compatible with the CHARMM all-atom additive biological force fields, J. Comput. Chem. 31 (2009) NA–NA. https://doi.org/10.1002/jcc.21367.

[54] S. Miyamoto, P.A. Kollman, Settle: An analytical version of the SHAKE and RATTLE algorithm for rigid water models, J. Comput. Chem. 13 (1992) 952–962. https://doi.org/10.1002/jcc.540130805.

[55] B. Hess, H. Bekker, H.J.C. Berendsen, J.G.E.M. Fraaije, LINCS: A linear constraint solver for molecular simulations, J. Comput. Chem. 18 (1997) 1463–1472. https://doi.org/10.1002/(SICI)1096-987X(199709)18:12<1463::AID-JCC4>3.0.CO;2-H.

[56] T. Darden, D. York, L. Pedersen, Particle mesh Ewald: An N·log(N) method for Ewald sums in large systems, J. Chem. Phys. 98 (1993) 10089. https://doi.org/10.1063/1.464397.

